# Adaptive and pleiotropic effects of evolution in synonymous sugar environments

**DOI:** 10.1101/2024.01.28.577607

**Authors:** Neetika Ahlawat, Pavithra Venkataraman, Raman Gulab Brajesh, Supreet Saini

## Abstract

Adaptation to an environment is enabled by the accumulation of beneficial mutations. When adapted populations are shifted to other environments, the byproduct or pleiotropic fitness effects of these mutations can be wide-ranged. Since there exists no molecular framework to quantify relatedness of environments, predicting pleiotropic effects based on adaptation has been challenging. In this work, we ask if evolution in highly similar environments elicits correlated adaptive and pleiotropic responses. We evolve replicate populations of *Escherichia coli* in non-stressful environments that contain either a mixture of glucose and galactose, lactose, or melibiose as the source of carbon. We term these similar sugars as “synonymous”, since lactose and melibiose are disaccharides made up of glucose and galactose. Therefore, the evolution environments differed only in the way carbon was presented to the bacterial population. After 300 generations of evolution, we see that the adaptive responses of these populations are not predictable. We investigate the pleiotropic effects of adaptation in a range of non-synonymous environments, and show that despite uncorrelated adaptive changes, the nature of pleiotropic effects is largely predictable based on the fitness of the ancestor in the non-home environments. Overall, our results highlight how subtle changes in the environment can alter adaptation, but despite sequence-level variations, pleiotropy is qualitatively predictable.

**Lay Summary:** In nature, evolution in “similar” environments is believed to elicit identical responses. For example, the arctic fox and ptarmigan, which are two unrelated species living in the arctic, have evolved to turn white in the winters. They did not evolve this ability because they from the common ancestor, but because the environment favoured this trait. In this work, we ask what happens to evolving populations if there are minute changes in the environment, and what are the consequences of adapting in these environments that are “almost identical”, or as we call them, “synonymous”.

We evolve replicate populations of the bacteria *E. coli* in three synonymous environments, and quantify their ability to grow in both synonymous and non-synonymous environments. We see that evolution does not proceed in an identical fashion in these populations, and that each environment favours a different trait. However, interestingly, in non-synonymous environments, these three sets of populations perform almost identically, and their growth is qualitatively predictable.

Our results show that even simple and subtle changes in the environment can act as drivers of biodiversity.

## Introduction

An evolving population accumulates mutations, and selection increases the frequency of beneficial mutations in the population. Mutational effects are contingent on the genetic background on which they occur (Bateson et al., 1909), and the environment that the population evolves in (Plate, 1910). As a result, adaptation in one environment could lead to pleiotropic effects in the others. Pleiotropy was thought to operate only antagonistically (Levins, 1974; Stearns, 1989). However, many ecological and laboratory studies have shown the evolution of generalists, which are fit in a wide-range of environments (Bedhomme et al., 2015; Forister et al., 2012; Geiler-Samerotte et al., 2020; Hall et al., 2006; Kassen, 2002; C. E. Lee, 2002; Mitchell-Olds et al., 2007; Savolainen et al., 2013; Schluter, 2009; Wang et al., 2015). Thus, the potential effects of pleiotropy are important, especially to understand processes such as ecological specialization (Friberg et al., 2008; Hardy & Forister, 2023; Ostrowski et al., 2007), genetic diversification (D.-C. Lee et al., 2023; Noda-Garcia et al., 2019; Solovieff et al., 2013), and movement on a rugged fitness landscape (W. Huang et al., 2012; Martin & Lenormand, 2006). Pleiotropic alleles have also been implicated in a range of human diseases (Alzheimer’s Disease Genetics Consortium et al., 2018; Bellou et al., 2020; Byars & Voskarides, 2020; Sivakumaran et al., 2011), and are known to confer antibiotic resistance (Hershberg, 2017; Pietsch et al., 2017; Schenk et al., 2015; Xia et al., 2017).

Several laboratory studies have focused on understanding the dynamics and variability of the pleiotropic effects of adaptation (Bailey & Kassen, 2012; Bakerlee et al., 2021; Cooper & Lenski, 2000; Dillon et al., 2016; Jasmin et al., 2012; Jerison et al., 2020; Kinsler et al., 2020; Leiby & Marx, 2014; Meyer et al., 2010; Novak et al., 2006; Ostrowski et al., 2005). Two types of populations have been seen to evolve in these experiments – generalists and specialists. Generalists are populations that are viable in a range of environments, whereas specialists incur significant fitness costs in non-home environments. While the evolution of specialists could be due to antagonistic pleiotropy, generalists could evolve due to mutations in genes that affect a small set of phenotypes (Crozat et al., 2010; Fumasoni & Murray, 2020; Good et al., 2017; C.-J. Huang et al., 2018; Kinsler et al., 2020; Rodríguez-Verdugo et al., 2013; Tenaillon et al., 2012; Venkataram et al., 2016). The fitness effect of a mutation in an environment which resembles the evolution environment is expected to be similar, because of relatedness in the growth conditions. For example, Bakerlee et. al. show that yeast populations that adapted in temperature stress showed increased fitness in a highly saline environment (Bakerlee et al., 2021). However, the same population’s fitness is much lower when tested in an environment colder than the evolution environment. Clearly, due to the lack of generality, adaptation to a stress does not guarantee an identical change in fitness when exposed to other stresses.

It is therefore not well understood how pleiotropic effects change with environment. In other words, we do not know what kind of environments drive the evolution of generalists or specialists. How do subtle changes in the evolution environment alter adaptation and pleiotropic effects? Is the nature and number of genetic changes in generalists predictable? And most importantly, how general is the behaviour of generalists?

In this study, we propose to answer these questions by carrying out laboratory evolution using *Escherichia coli* in three evolution environments (Supplement Figure 1). The environments differ in the source of carbon available for the microbes to feed on. The three evolution environments contained one of the following sugars - (a) a mixture of glucose and galactose, (b) lactose, (c) melibiose. Glucose and galactose, when linked by a β-1,4-glycosidic and an α-1,6-glucosidic bond form lactose and melibiose respectively. Since these three sugars are different combinations of glucose and galactose, we term them as “synonymous” environments. Essentially, the evolution environments differed only in the way glucose and galactose were presented to the evolving population.

We evolved six replicate populations in each of the synonymous environments for 300 generations, and assayed the fitness of the evolved populations in synonymous and non-synonymous sugar environments. Our results allow us to identify the pleiotropic effects of adaptation in a not-so-harsh environment, and subsequently comment on the predictability of the evolution of specialists/generalists. Most importantly, in a novel attempt, we show that minute changes in resource packaging can alter adaptation and the consequent pleiotropic effects.

## Results

### Adaptation in synonymous sugars is non-identical

Replicate populations of *E. coli* were evolved in M9 minimal media containing 0.2% of glucose-galactose (0.1% of each), lactose, or melibiose. After 300 generations, two fitness parameters – growth rate in the exponential phase and carrying capacity – were calculated.

As shown in **Figure 1**, the biomass accumulated at the end of 16 hours increased for all the populations relative to the ancestor (*p<0.05, one-tailed t-test*), and other than two melibiose-evolved cells, growth rate increased in all cases (*p<0.05, one-tailed t-test*). Interestingly, two out of the six melibiose-evolved populations’ growth rates are lower than that of the ancestor (*p<0.05, one-tailed t-test*).

**Figure 1.**
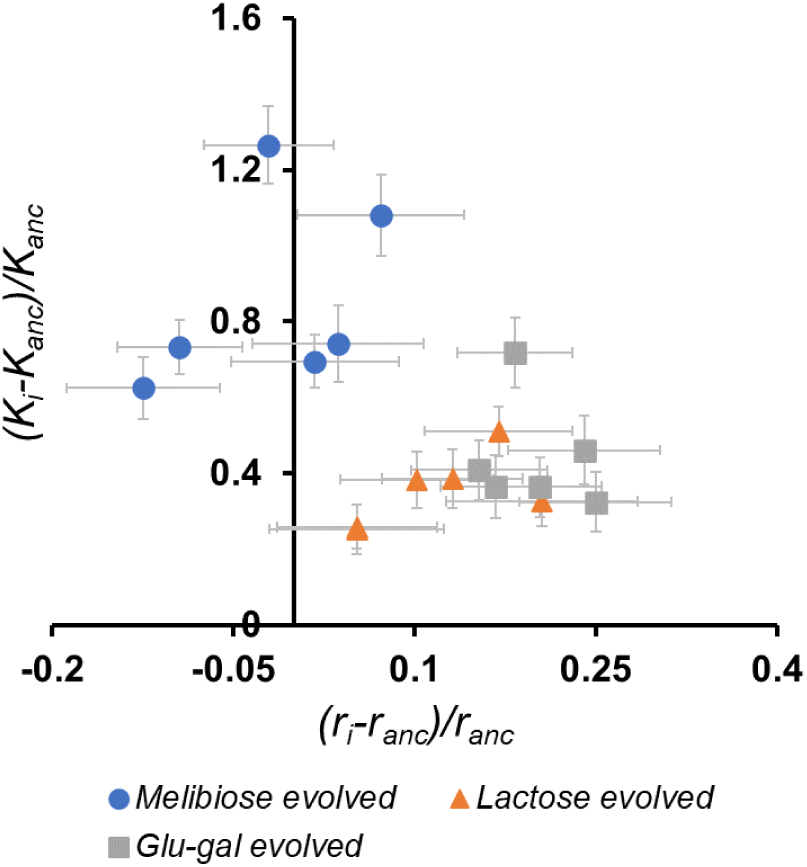
Selection acts on different traits in different evolution environments. Six replicate populations of *E. coli,* founded from a single clone, were evolved in three different, yet synonymous sugar environments – glucose-galactose mixture, lactose, and melibiose. After 300 generations of evolution, fitness assays show that the growth rates (*r*) and carrying capacities (*K*) of almost all the evolved populations increased relative to the ancestor. However, the melibiose-evolved populations showed a much larger increase in carrying capacity than the other two sets of populations. In fact, two of the melibiose-evolved populations’ growth rates decreased relative to the ancestor. Therefore, our results show that selection acting on a population changes even with a minor change in the environment.

Our results show that the selection pressure imposed by synonymous environments is non-identical. In fact, the melibiose evolved populations show a qualitative change in adaptive response compared to the lactose and glucose-galactose evolved populations, whose fitness differ from each other only quantitatively.

Populations with an increased growth rate (*r*) are fast growers, while populations with an increased carrying capacity (*K*) are considered better competitors (Pianka, 1970). While intuition suggests that evolving populations should exhibit *r-K* trade-offs, there exists very little empirical evidence in support. Recently, Marshall et al. showed theoretically how and why *r* and *K* may covary positively (Marshall et al., 2023). However, Wei et al. show that reduction in environment quality leads to such “trade-ups” in *r* and *K* (Wei & Zhang, 2019). Our results show that trade-ups in *r* and *K* are commonly observed in populations that evolved in different sources of carbon, and that the rate of occurrences of trade-ups or trade-offs is dependent on the exact nature of the evolution environment.

### Pleiotropic effects of adaptation in synonymous environments

Next, we shift these three sets of evolved populations to non-home environments, that contain synonymous sugars as the source of carbon. We calculate the growth *r* and *K* at the end of 16 hours, assuming exponential growth. Fitness changes of twelve populations in non-home synonymous environments, relative to the ancestor, are as shown in **Figure 2a** and **2b**.

**Figure 2.**
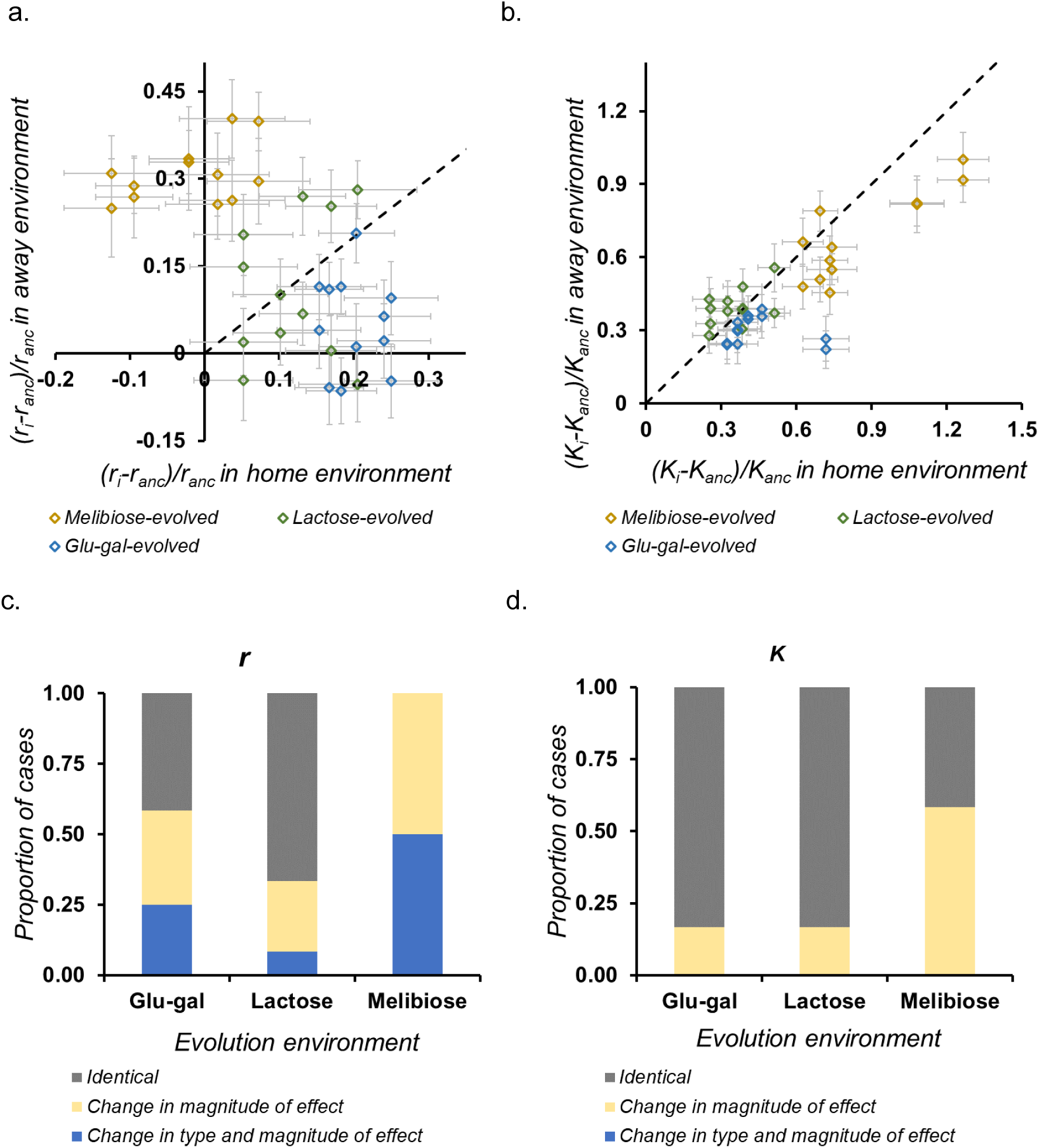
Pleiotropic responses of evolved populations in synonymous environments. We tested the fitness (*r* and *K*) of the three sets of evolved populations in away environments, in which the source of carbon was a synonymous sugar, and checked if fitness gains in home and away environments were correlated. For each set of evolved population, there were two synonymous away environments in our experiment. **(a)** and **(b)** show the relative gains in home and away environments, in *r* and *K,* respectively. We count the number of distinct types of pleiotropic effects. As shown in **(c)** and **(d)**, the relative frequencies of the three types of pleiotropic responses are different for different sets of evolved populations.

Next, the fitness gains for each population relative to the ancestor, in home and non-home environments were compared. Each such comparison can be classified into one of the following three categories: (a) identical to the ancestor, in magnitude and type of effect, (b) identical to the ancestor, in the type of effect but is different in magnitude, or (c) different from the ancestor, in the type and magnitude of effect. **Figure 2c & 2d** shows the proportion of these cases (out of 12), for *r* and *K*, for the three sets of evolved populations, when grown in non-home synonymous environments.

Correlated fitness gains are the most common in the lactose-evolved populations, followed by the glucose-galactose-evolved populations. The melibiose-evolved populations, show the most diverse range of pleiotropic responses – growth rates of all the melibiose-evolved populations were non-identical in the synonymous environments, and the carrying capacity of more than half of them was different from that in the home environment.

By and large, our results agree with the observations made by Ostrowski et. al., that adaptation in a sugar environment confers positive pleiotropic effects in other sugars (Ostrowski et al., 2005). However, despite being shifted to synonymous environments, our evolved populations show considerable variability in their pleiotropic responses. What happens to variability in pleiotropic responses in non-synonymous sugars?

### Pleiotropic effects of adaptation in non-synonymous environments

The three sets of populations which evolved independently in melibiose, lactose, and glucose-galactose were shifted to non-synonymous sugar environments and tested for fitness. These non-synonymous environments consisted of one of pentoses (arabinose, xylose), a methyl pentose (rhamnose), sugar alcohols (glycerol, sorbitol), hexoses (fructose, mannose), or a trisaccharide (raffinose) as the source of carbon. Again, we measured *r* in exponential phase and *K* as cells enter stationary phase. Fitness changes of the eighteen evolved populations, relative to that of the ancestor, are shown in **Figure 3a** and **3b**. We found no environment in which the fitness of all the evolved populations dropped compared to the ancestor. Arabinose and sorbitol were two environments in which both *r* and *K* of all the evolved populations increased relative to the ancestor. Therefore, all the populations that evolved in one of the synonymous environments were fit in arabinose and sorbitol. However, we did not find any set of evolved populations in which all the replicates performed identically in home and any of the non-home environments. As a result, given a non-home environment, we could only say if an evolved population is fitter than the ancestor or not; we were unable to comment on the exact difference in fitness based on the fitness of the evolved population in the home environment.

**Figure 3.**
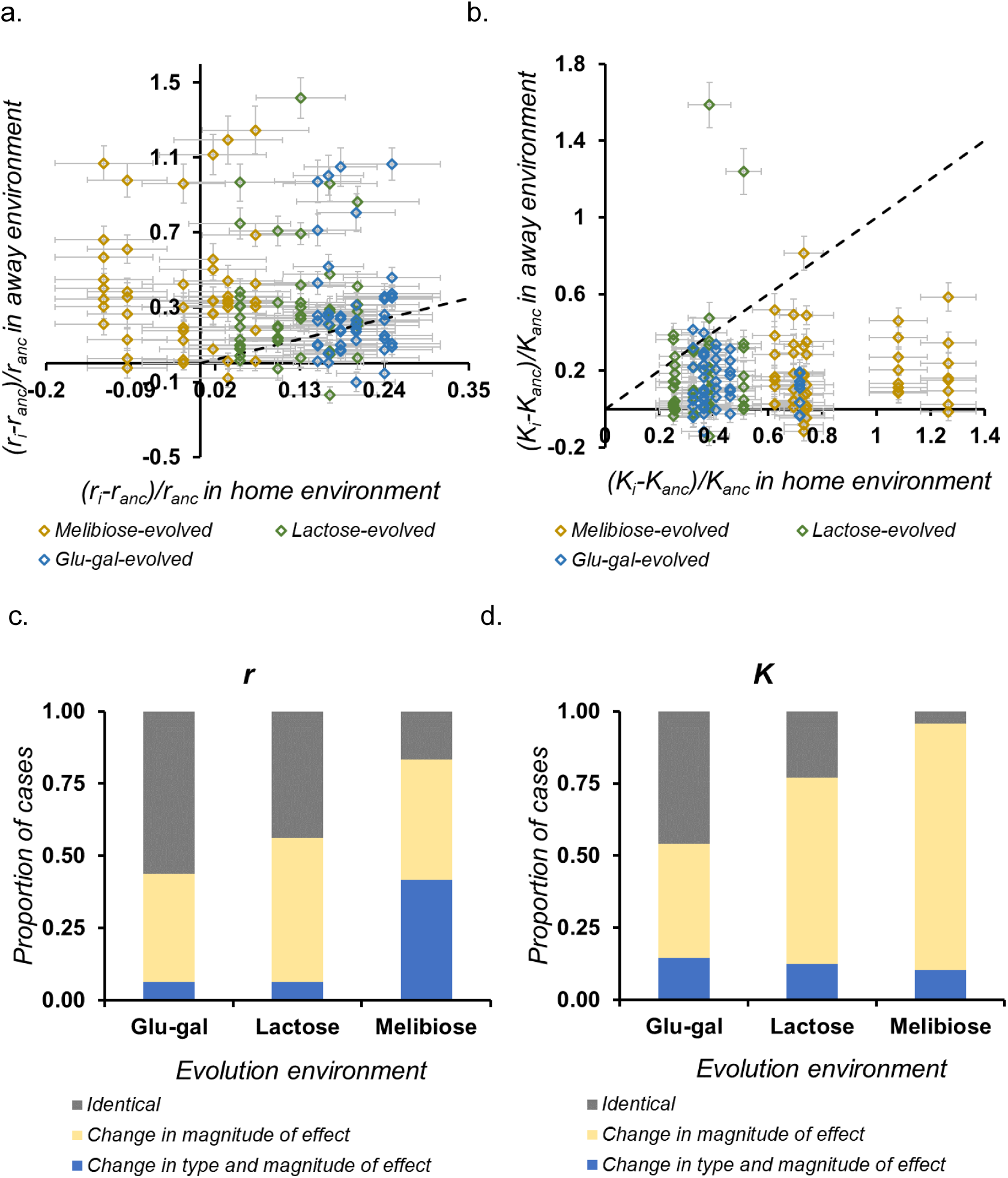
Pleiotropic responses of evolved populations in non-synonymous environments. We tested the fitness (*r* and *K*) of the three sets of evolved populations in away environments, in which the source of carbon was a non-synonymous sugar, and checked if fitness gains in home and away environments were correlated. For each set of evolved population, there were eight non-synonymous away environments in our experiment. **(a)** and **(b)** show the relative gains in home and away environments, in *r* and *K,* respectively. We count the number of distinct types of pleiotropic effects, and the results are as shown in **(c)** and **(d)**.

In the eight non-synonymous sources of carbon, glucose-galactose evolved populations showed the maximum number of similar fitness changes, followed by the lactose-evolved populations. The melibiose-evolved populations continued to show diverse responses in non-synonymous environments as well, as shown in **Figure 3c and 3d**.

**Figure 4** shows the overall proportion of cases (considering both *r* and *K*, and synonymous and non-synonymous environments) of pleiotropy for the three sets of evolved populations. It is evident that the variability in the type of pleiotropic responses depends on their evolution environment.

**Figure 4.**
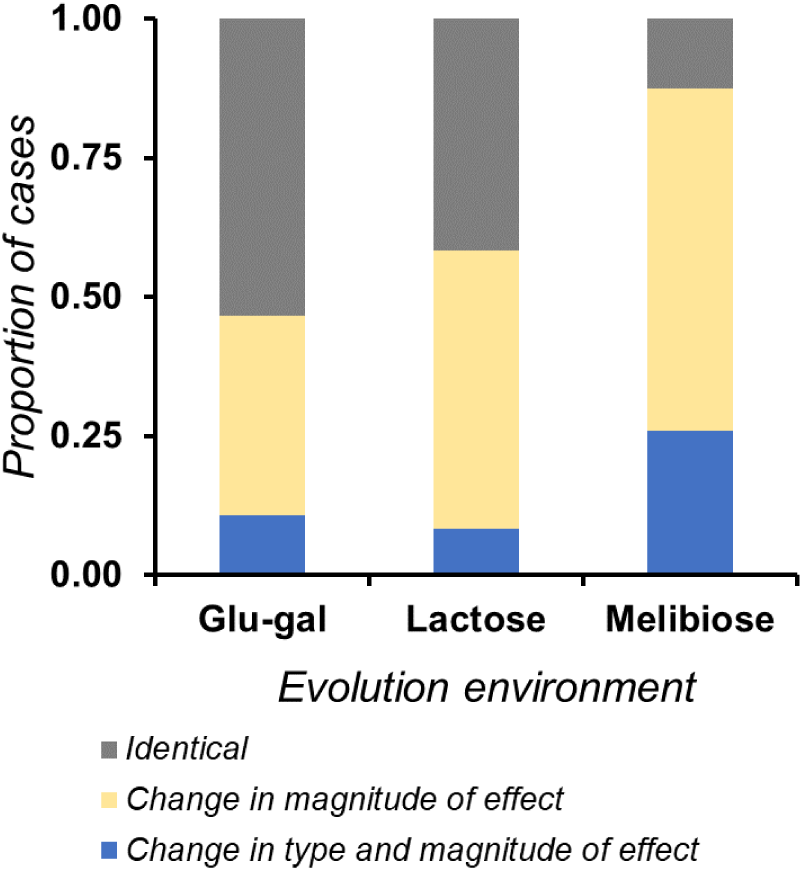
Variability in pleiotropic responses. To test whether the pleiotropic fitness changes (both *r* and *K*) of glucose-galactose, lactose, and melibiose evolved populations in away environments (both synonymous and non-synonymous) follow any patterns, we compared the relative frequencies of the three types of pleiotropic responses exhibited by the three sets of evolved populations. Melibiose evolved populations exhibited the greatest number of pleiotropic responses, followed by the lactose evolved populations. Overall, we observe that even minute changes in the evolution environment play a significant role in dictating the variability in pleiotropic responses.

### Fitness of the ancestor as a predictor of pleiotropic effects

Global epistasis has enabled the prediction of the fitness effect of a beneficial mutation as a function of the fitness of the background on which it occurs (Kryazhimskiy et al., 2014). We tried to identify if ancestral fitness can help predict pleiotropic effects. Specifically, we checked if the pleiotropic fitness of an evolved population correlated with the fitness of the ancestral population in any given environment. Therefore, we compared the fitness changes in the evolved populations with that of the ancestor, in both synonymous and non-synonymous environments.

In synonymous environments, there existed no correlation between ancestral fitness and the nature of pleiotropic responses, as shown in **Figure 5a and 5b**. The three sets of evolved populations showed qualitatively similar pleiotropic responses in the non-synonymous environments (refer to Supplement Figures 2 & 3). As shown in **Figures 5c and 5d**, in non-synonymous environments, average gains in *r* and *K* correlated negatively with ancestral fitness.

**Figure 5.**
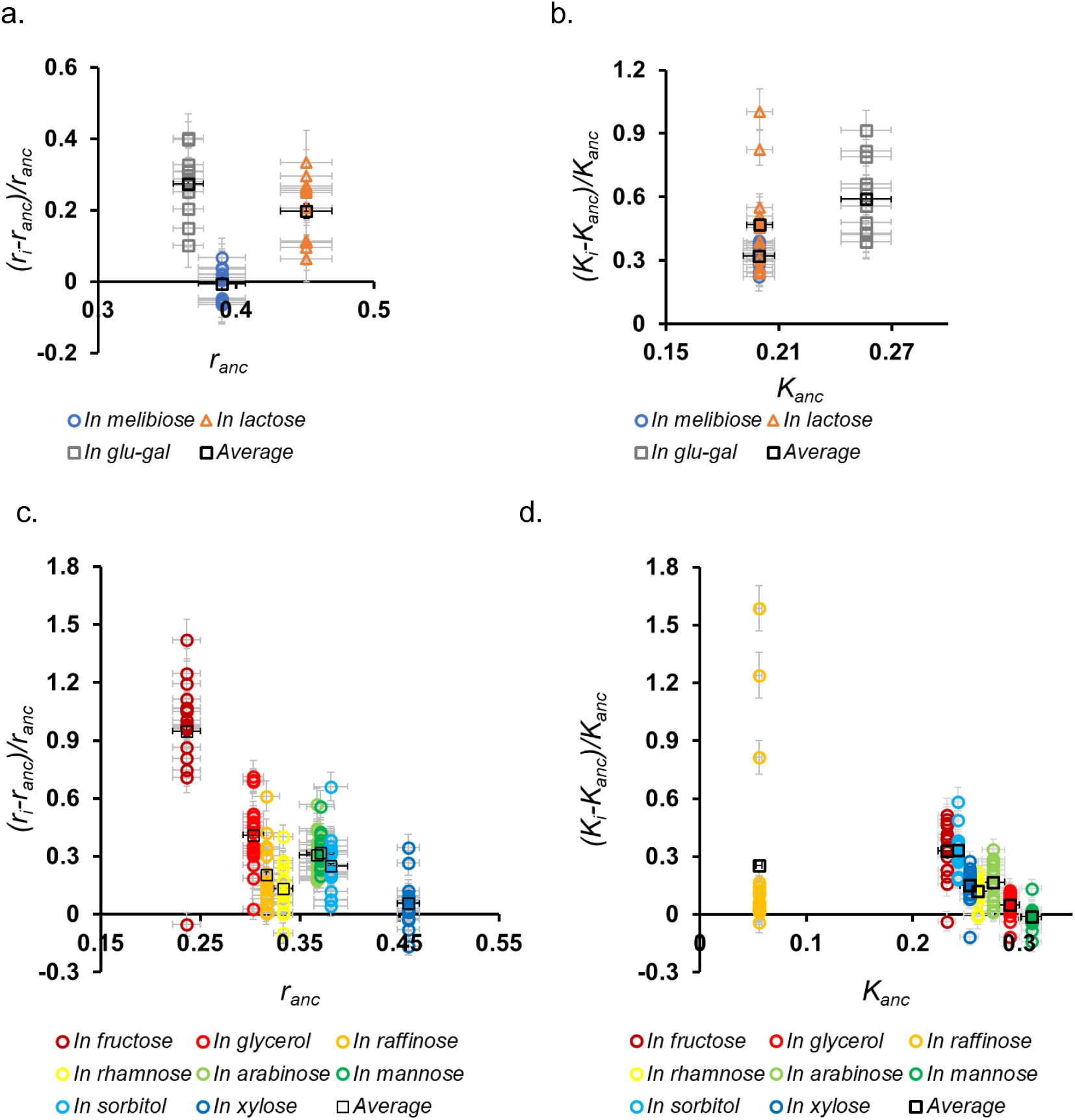
Fitness of the ancestor could act as a predictor of pleiotropic fitness gains. We test if the fitness of the ancestral population in an environment can act as a predictor of the fitness of an evolved population. We compare the fitness gain of an evolved population with ancestral fitness in away environments. **(a)** and **(b)** show these comparisons for *r* and *K* in synonymous environments, respectively. It must be noted that for each set of evolved populations, there were two synonymous away environments. We also plot relative fitness gains against ancestral fitness in eight non-synonymous environments. As shown in **(c)** and **(d)**, in most cases, there exists a trend – in non-synonymous environments, average pleiotropic fitness gains decrease with an increase in the fitness of the ancestor. However, there exists no such quantitative trend in the case of synonymous environments.

We next check if the variability in pleiotropic responses of these evolved populations is predictable based on ancestor’s fitness in the non-home environment. To do so, we consider the range of fitness gains (in *r* and *K*), calculated as the difference between the maximum and minimum of the mean fitness gain, as a proxy for variability in pleiotropic responses. As shown in **Figure 6a** and **6b**, as the fitness of the ancestor increases in the non-home environment, the variability in pleiotropic responses decreases in most cases. But variability did not correlate with average fitness changes. Therefore, given the fitness of the ancestor in a non-home environment, we show that a qualitative prediction of pleiotropic effects of an evolved population is possible.

**Figure 6.**
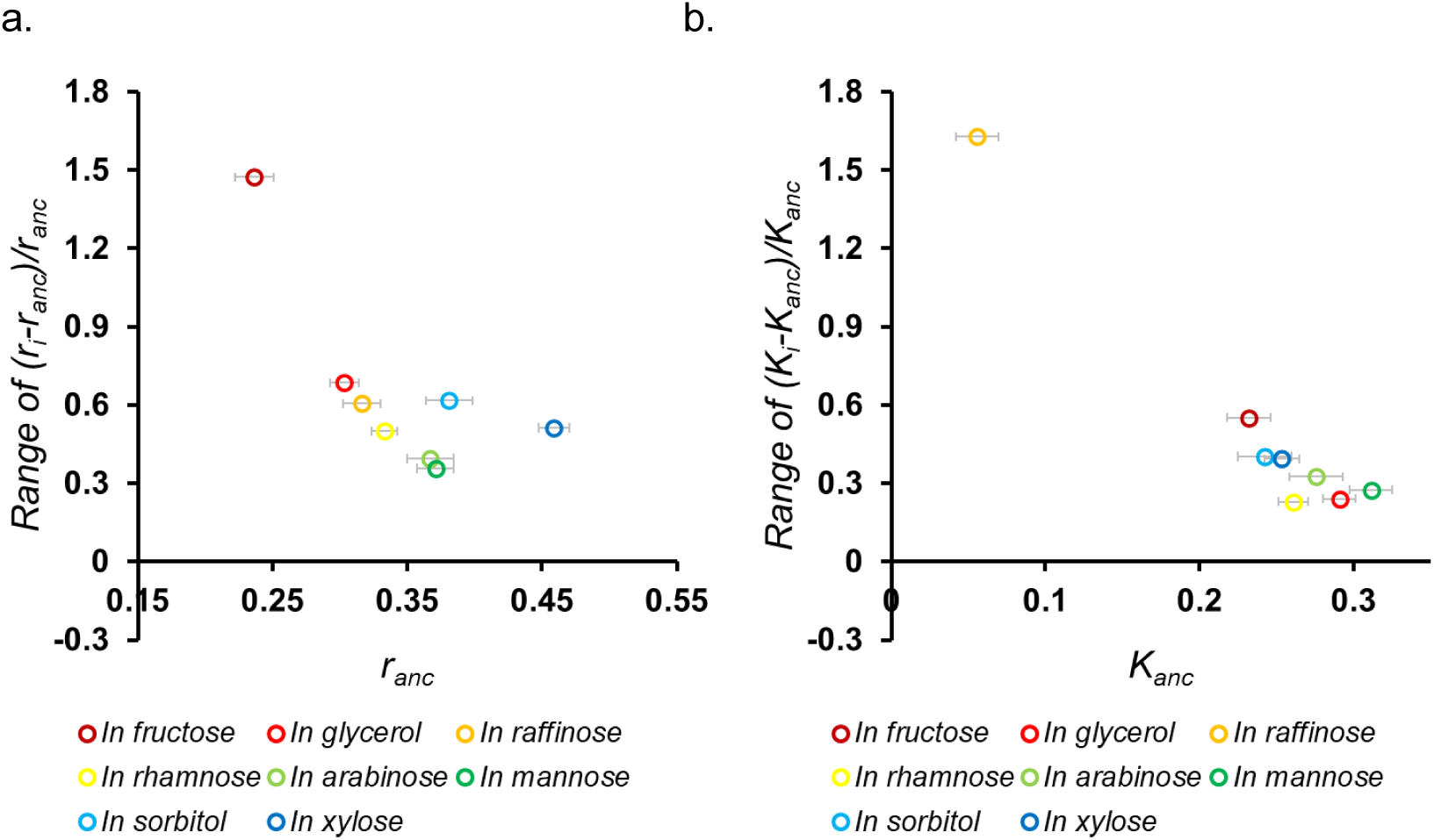
Fitness of the ancestor could act as a predictor of the variability in pleiotropic fitness gains. We showed that the fitness of the ancestor could as a rough predictor of the average pleiotropic fitness gain. Next, to test if the variability in pleiotropic responses can also be predicted in a similar fashion, we plot the range of fitness gains in *r* (as shown in **(a)**) and *K* (as shown in **(b)**) against ancestral fitness in the eight non-synonymous environments. Range was calculated as the difference between the maximum and minimum fitness gains in the eighteen evolved populations.Variability in pleiotropic responses, like mean pleiotropic fitness gain, also decreases with increase in ancestor’s fitness, in most cases.

### The underlying genetic diversity explains the nature of pleiotropic responses

We genome sequenced the eighteen evolved populations to identify the genetic changes accumulated in the three synonymous evolution environments. Mutation analyses reveal that mutations in the *rpo* genes were the most common among all sets of evolved populations. *Rpo* genes, that code for proteins that make up RNA polymerase, are known to facilitate adaptation in thermal, osmotic, and antibiotic stresses (Hews et al., 2019). Other mutational targets included *yafW* which is a part of a toxin-antitoxin pair, *nanR* and *acrR*, which are transcription regulators (Amores et al., 2017; Chu et al., 2008), and the methionine transporter *metN* (Kadaba et al., 2008). Overall, the melibiose-evolved populations had the highest number of distinct mutational targets, as shown in Figure 7a (Supplement Table 1 lists the mutations in each population). Figure 7b shows the correlated between the proportion of pleiotropic responses and the number of mutational targets.

**Figure 7.**
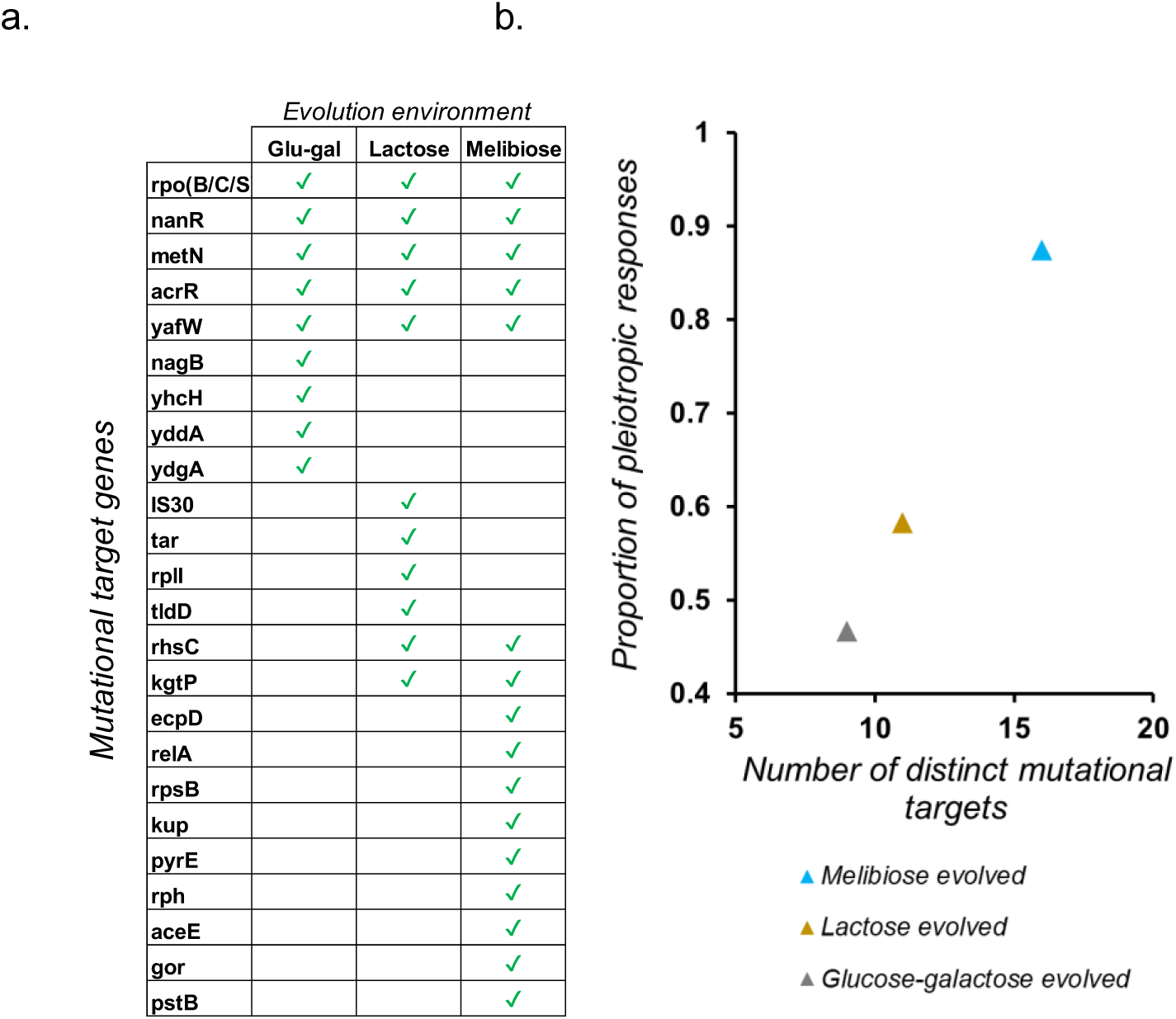
Genetic basis of pleiotropic effects. (a) The mutational targets in the three sets of evolved populations are shown. (b) The number of distinct mutational targets correlates with the proportion of pleiotropic responses. In other words, our results show that with an increase in the number of distinct types of mutational targets, proportion of pleiotropic effects of the evolved populations increases.

We counted the number of instances where the three sets of evolved populations showed any type of pleiotropy (change in magnitude and/or sign of fitness effect), across the ten non-home environments (two synonymous, eight non-synonymous) their fitness was measured in. Considering changes in both *r* and *K*, we see that pleiotropic effects are the maximum for the melibiose-evolved populations, as shown in Figure 7b. This proportion of pleiotropic responses, along with that of lactose-evolved and glucose-galactose evolved populations, correlated predictably with the number of distinct genes in which adaptive mutations occurred in replicate populations.

### Conclusions and Discussion

Pleiotropic consequences of adaptation are expected to vary with changes in the duration of evolution (Bailey & Kassen, 2012; Bakerlee et al., 2021; Cooper & Lenski, 2000; Meyer et al., 2010; Novak et al., 2006). It has been shown that evolving in a particular environment for a long duration leads to the rise of a specialist population (Bakerlee et al., 2021). However, the role of the environment in dictating the evolution of a particular type of population - specialists or generalists - is not well understood. Also, there exists no method to quantify the relatedness of environments, making the prediction of pleiotropy challenging.

To probe the influence of environmental factors in dictating evolutionary trajectories and pleiotropic effects, we evolve six replicate populations of *E. coli* for a short period of time (300 generations) to capture the early onset of pleiotropy. The evolution environments were not harsh – the ancestral cell was viable in the three sugar environments that we picked. The evolution of *E. coli* in glycerol and lactate (propane-based organic compounds) has been shown to yield identical phenotypic response (Fong et al., 2005). How widely applicable is this result? In this study, we presented to the population two monosaccharides (glucose and galactose) packaged in three different ways – a mixture of glucose and galactose, lactose, and melibiose. These “synonymous” sugar environments allow us to maintain similarity in the evolution environments.

In a novel attempt, we show that evolution in synonymous sugars does not elicit identical adaptive responses. It must be noted that lactose and melibiose are utilized almost identically by *E. coli –* they are first imported into the cell with the help of a permease, a galactosidase then breaks them down into glucose and galactose, which are further metabolized (Jacob & Monod, 1961; Pardee, 1957). Despite this similarity, we see that the pressure to accumulate mutations that improve biomass is the strongest in the presence of melibiose. In fact, in some cases, evolution in melibiose led to a decrease in growth rate. This is one of the few cases where a trade-off between *K* and *r* has been empirically reported (Beardmore et al., 2011; Fitzsimmons et al., 2010; Meyer et al., 2015; Novak et al., 2006; Reding-Roman et al., 2017; Warringer et al., 2011; Wei & Zhang, 2019). Commonly, growth rate is used as a proxy for fitness. Our observations underscore the need to look at other traits of evolving populations on which selection could be acting.

Fluctuating environments are known to generate genetic diversity (Abdul-Rahman et al., 2021; Ellegren & Galtier, 2016; Yamamichi et al., 2023), and alter gene expression via mutations in global regulators (López-Maury et al., 2008; Mahilkar et al., 2021; Podrabsky & Somero, 2004). By pointing at how minute changes in the evolution environment can also generate diverse genotypic changes, we have identified a simple yet significant driver of the process of generation of biodiversity in prokaryotes.

We expected evolution in synonymous sugars to elicit positive pleiotropic responses in synonymous environments (Jerison et al., 2020). Our fitness assays indicate clearly that positive pleiotropic fitness changes are common, in both synonymous and non-synonymous environments. In fact, we also find cases where the fitness effect in a non-home environment is greater than that in the home environment. For example, the growth rates of two melibiose-evolved populations decreased compared to the ancestor, in the home environment. However, both these populations showed an increase in growth rate in almost all the non-home environments. Interestingly, there were scenarios were populations that adapted in a synonymous environment *B* performed better in *A* than any of the populations that adapted in *A.* Therefore, moving to a not-so-different foreign environment could enable access to a fitness peak inaccessible to an evolving population.

Almost all the melibiose-evolved populations show considerable fitness gains in other synonymous and non-synonymous environments. However, why did these mutations not fix in the other synonymous environments? Recently, Kosterlitz *et. al.* reported that shifting hosts may aid adaptation of a gene, which are blocked due to epistatic interactions in the native host (Kosterlitz et al., 2023). Our results point at similar roadblocks to adaptation posed by the environment, and indicate how minute changes in the environment can help overcome them. It remains to be seen how the adaptive trajectories of these evolved populations differ, when allowed to evolve in a non-home environment for a long period of time.

The diversity in pleiotropic responses depended both on the home and the non-home environment. For example, in glycerol, all the eighteen evolved populations accumulated biomass different from that of the ancestor. In seven out of the eight non-synonymous sugars that we tested the fitness of the evolved populations in, over 50% of the populations showed pleiotropic effects.

Across all sets, we found more generalists than specialists at the end of our evolution experiment that lasted 300 generations. We attempted to identify if ancestral fitness can serve as an indicator of the exact pleiotropic effect. Surprisingly, we found no correlation of pleiotropic responses with ancestor fitness in synonymous environments, but there exists a pattern in non-synonymous environments. Much like how the theory of diminishing returns explains the fitness effect of new beneficial mutations given the background fitness (Diaz-Colunga et al., 2023), we show that given the fitness of the ancestor in an environment, it is possible to comment on the qualitative nature of the pleiotropic fitness of an evolved population despite variations and stochasticity at the genetic level brought about by adaptation. The number of instances where pleiotropic effects were observed (in non-home environments), whether antagonistic or not, was understandably correlated with the number of distinct mutational targets, as reported in the past (Kinsler et al., 2020). Adaptive mutations occurred in global regulators like the *rpo* genes, but different SNPs in a gene elicited variable adaptive and pleiotropic effects. Therefore, to quantify the exact pleiotropic fitness effects, our results show that a sequence level mapping of fitness effects and environments is necessary.

Studies of pleiotropy have largely focused on studying the effects of evolving in one stress environment in another. Our experimental approach provides a framework to study evolution and its consequences in related environments, while evolving the microbial population in a non-stress environment. Our attempt, in this work, has been focused on being able to quantify and predict pleiotropy in related environments (sugar environments), and not in highly dissimilar environments. We aim to identify the byproduct effects of these adapted populations in unrelated environments (temperature stress, osmotic stress, etc.) in the future, and test how changing the qualitative nature of the environment alters pleiotropic responses. It is also unclear how the trends and correlations we observe will change with time, and if and when the generalist populations will become specialists. Therefore, we continue to propagate these bacterial populations in their respective environments for future studies.

Overall, our work motivates further probe into the temporal variability of adaptation in synonymous environments. The results we report show that global epistasis renders ancestral fitness as a predictor for early effects of pleiotropy, and point at the need to perform high-throughput evolution experiments to build a framework that can help decipher pleiotropic effects in a range of non-home environments.

## Material and Methods

### Strains used and media composition

*E. coli* K12 MG1655 (ATCC 47076) was used in this study.

### Evolution experiments

The evolution experiment was conducted in M9 minimal media (composition: Na_2_HPO_4_.7H_2_O (12.8 g/L), KH_2_PO_4_ (3g/L), NaCl (0.5g/L), NH_4_Cl (1g/L), 1M MgSO_4_ and 0.1M CaCl_2_), containing either a mixture of glucose and galactose (0.1% each) or lactose (0.1%) or melibiose (0.1%).

Six independent replicate populations of *E. coli* were evolved at 37⁰C and at 250 rpm in three environments - glucose-galactose mixture, lactose, or melibiose. Evolution experiment was performed by serially transferring 50μL of culture into 4.95mL of fresh M9 media containing appropriate sugar, every 12 hours.

### Fitness measurements

To measure fitness, cells were grown in LB (1% tryptone, 0.5% yeast extract, 0.05% NaCl) and incubated at 37⁰C for 12 hours with shaking at 250 rpm. Thereafter, cells were sub-cultured 1:100 in fresh M9 media containing 0.2% glycerol for 18 hours. To measure fitness, cells were cultured in M9 media containing the appropriate carbon environment (glucose-galactose, lactose, melibiose, L-arabinose, D-xylose, D-mannose, D-fructose, D-sorbitol, D-raffinose) to an initial OD600 of 0.01.

Cells were then transferred to a 96-well plate (Costar) and the plate was covered with a breathable membrane (Breathe-Easy) to prevent evaporation. The 96-well plate was incubated at 37⁰C with shaking in a microplate reader (Tecan Infinite Pro 200). OD600 was measured every 30 minutes until each culture reached stationary phase (∼20 hours).

Growth rate was measured as described in the past (Choudhury & Saini, 2019). Maximum OD attained in each line was used as a measure for biomass accumulation. There was no observable lag phase in the glucose-galactose and lactose-evolved populations, as observed in another study (Choudhury & Saini, 2019). Therefore, we did not consider it as a measure of fitness change.

### Statistical tests

The error bars in the plots shown correspond to standard deviation, unless specified otherwise. A comparison of two means was done using a two-tailed *t* test. A one-tailed *t* test was used to ascertain if one mean is greater than or lesser than another. In all cases, the significance level was set to 0.05.

### Whole-genome sequencing

Genomic DNA of ancestral and evolved populations of *E. coli* was isolated using the standard phenol/chloroform DNA isolation protocol (He, 2011). DNA quality and concentration were measured after DNA isolation using Nanodrop Spectrophotometer (Eppendorf), and by gel electrophoresis. Genomic DNA samples were sent for paired-end sequencing using Illumina NovaSeq 6000, with an average read-depth of 151bp.

Based on the quality report of fastq files, sequences were trimmed to retain only high-quality sequences for analysis. The adapter trimmed reads were aligned to the reference genome, *Escherichia coli* (ATCC 47076) using Burrows Wheeler Aligner (BWA). Each sample had a minimum coverage of more than 30x. Variant calling was done for the samples using GATK and further annotated using SnpEff. Variants that were present in the ancestral strain were filtered out manually. The remaining SNPs were used for further analysis. Raw sequencing data is available at <u>ncbi.nlm.nih.gov/sra/PRJNA1022868</u>.

## Acknowledgements.

This work was funded by a DBT/Wellcome Trust (India Alliance) grant (Award No. IA/S/19/2/504632) to SS. PV is supported by the Prime Minister’s Research Fellowship (PMRF ID 1302050). NA is supported by the Institute Post-Doctoral Fellowship by IIT Bombay.

