## Supplement Figures and Tables for "Adaptive and pleiotropic effects of evolution in synonymous sugar environments"

### Supplementary Figures and Table.

#### Figure S1.

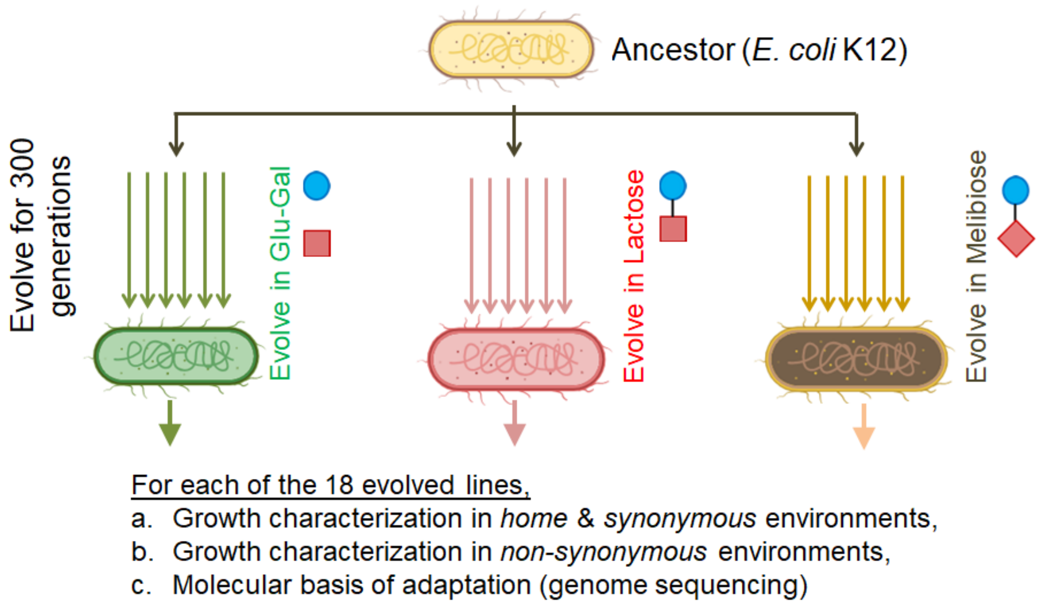


**Supplementary Figure 1. Experimental design of this study.** Ancestor *E. coli* was evolved in three 'synonymous' environments – a mixture of glucose (blue circle) & galactose (red square); lactose; and melibiose for approximately 300 generations. We evolved six independent lines in each of the three environments. After evolution via serial sub-culture (1:100 every 12 hours), the growth characteristics of the evolved lines were studied in *home, synonymous*, and *non-synonymous* environments. Genome sequencing was performed for the 18 evolved lines to identity the molecular basis of adaptation.

**Figure S1.**

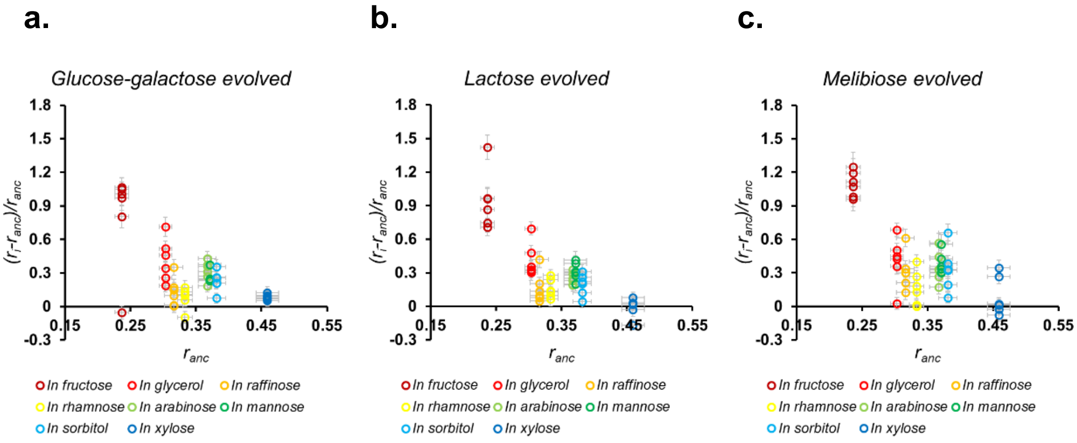


**Supplementary Figure 2. Growth rate of the ancestor as a predictor of pleiotropic growth rate.** We test if the growth rate of the ancestral population in a range of non-synonymous environments correlates with the fitness changes of the evolved populations. Glucose-galactose (shown in **(a)**), lactose (shown in **(b)**) and melibiose(shown in **(c)**) evolved populations show qualitatively similar pleiotropic fitness effects.

**Figure S3.**

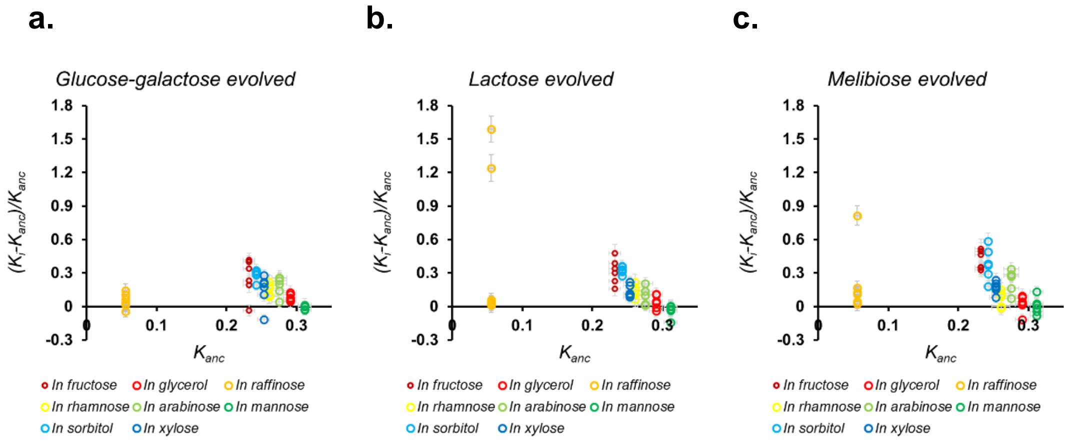


**Supplementary Figure 3. Biomass accumulation of the ancestor as a predictor of pleiotropic biomass accumulation.** We test if the biomass accumulated by the ancestral population in a range of non-synonymous environments correlates with the fitness changes of the evolved populations. Glucose-galactose (shown in **(a)**), lactose (shown in **(b)**) and melibiose(shown in **(c)**) evolved populations show qualitatively similar pleiotropic fitness effects.

**Supplementary Table 1. List of mutations in the evolved populations.** All the evolved populations were genome-sequenced to identify adaptive mutations. **(a)** shows the mutations in the six glucose-galactose evolved populations, **(b)** shows the mutations in the lactose evolved populations, and **(c)** shows the mutations in the melibiose evolved populations.

**a.**

| Evolved lines | Read depth | Gene name | Position of the variant in the gene (HGVS.c) |
|---|---|---|---|
| gG1 | 27 | acrR | c.-4819T>C |
| | 84 | yafW | c.150C>A |
| | 16 | metN | c.-1096C>T |
| | 117 | rpoC | c.1865A>G |
| | 35 | nanR_2 | c.-706C>T |
| gG2 | 15 | IS30 | c.781C>G |
| | 268 | YafW | c.150C>A |
| | 15 | ycgJ | c.-3237_-3236ins[T] |
| | 164 | rpoB | c.1576C>T |
| | 49 | ydgA_2 | c.-3671A>C |
| | 39 | nanR_2 | c.-706C>T |
| gG3 | 25 | yhcH_1 | c.204T>G |
| | 19 | acrR | c.-4819T>C |
| | 21 | nagB_1 | c.-4677C>G |
| | 21 | nagB_1 | c.-4678C>G |
| | 49 | yddA | c.-12C>T |
| | 118 | yafW | c.150C>A |
| | 108 | rpoC | c.1865A>T |
| | 31 | ydgA_2 | c.-3671A>C |
| | 25 | nanR_2 | c.-706C>T |
| gG4 | 33 | acrR | c.-4819T>C |
| | 68 | yafW | c.150C>A |
| | 114 | rpoC | c.1865A>G |
| | 23 | ydgA_2 | c.-3671A>C |
| | 17 | nanR_2 | c.-706C>T |
| gG5 | 19 | rcnB | c.-1531T>G |
| | 80 | yafW | c.150C>A |
| | 15 | metN | c.-1096C>T |
| | 44 | rpoC | c.1865A>G |
| | 19 | nanR_2 | c.-706C>T |
| gG6 | 89 | yafW | c.150C>A |
| | 123 | rpoB | c.1576C>T |
| | 22 | nanR_2 | c.-706C>T |

**b.**

| Evolved lines | Read depth | Gene name | Position of the variant in the gene (HGVS.c) |
|---|---|---|---|
| L1 | 42 | acrR | c.-4819T>C |
| | 130 | rpoS | c.361G>A |
| | 22 | kgtP | c.-431C>A |
| | 22 | rhsC_3 | c.724T>G |
| | 196 | yafW | c.150C>A |
| | 27 | ycgJ | c.-3237_-3236ins[T] |
| | 33 | metN | c.-1096C>T |
| | 133 | rpoC | c.3593T>A |
| | 18 | ydgA_2 | c.-3671A>C |
| | 22 | ydgA_2 | c.-3763C>G |
| | 46 | nanR_2 | c.-706C>T |
| L2 | 103 | rpoS | c.544G>C |
| | 68 | yafW | c.150C>A |
| | 130 | rpoB | c.1576C>T |
| | 16 | nanR_2 | c.-706C>T |
| L3 | 21 | acrR | c.-4819T>C |
| | 164 | rpoS | c.544G>C |
| | 146 | yafW | c.150C>A |
| | 17 | metN | c.-1096C>T |
| | 58 | rplI | c.309_312del[AACT] |
| | 41 | nanR_2 | c.-706C>T |
| L4 | 19 | acrR | c.-4819T>C |
| | 26 | tldD | c.622T>G |
| | 82 | yafW | c.150C>A |
| | 21 | ycgJ | c.-3237_-3236ins[T] |
| | 18 | metN | c.-1096C>T |
| | 137 | rpoB | c.1576C>T |
| | 16 | nanR_2 | c.-706C>T |
| L5 | 26 | acrR | c.-4819T>C |
| | 28 | tldD | c.622T>G |
| | 15 | kgtP | c.-431C>A |
| | 69 | yafW | c.150C>A |
| | 18 | metN | c.-1096C>T |
| | 123 | rpoB | c.1576C>T |
| | 21 | nanR_2 | c.-706C>T |
| L6 | 19 | acrR | c.-4819T>C |
| | 47 | rpoS | c.361G>A |
| | 31 | tar | c.1467G>T |
| | 24 | IS30 | c.18C>A |

**c.**

| Evolved lines | Read depth | Gene name | Position of the variant in the gene (HGVS.c) |
|---|---|---|---|
| M1 | 117 | rpoS | c.544G>C |
| | 15 | kgtP | c.-431C>A |
| | 36 | ecpD | c.36C>T |
| | 108 | yafW | c.150C>A |
| | 26 | metN | c.-1096C>T |
| | 19 | nanR_2 | c.-706C>T |
| M2 | 64 | yafW | c.150C>A |
| | 16 | metN | c.-1096C>T |
| | 20 | nanR_2 | c.-706C>T |
| M3 | 34 | relA | c.664C>A |
| | 67 | yafW | c.150C>A |
| | 97 | rpsB | c.684dup[T] |
| | 16 | kup | c.-4215G>T |
| M4 | 27 | rph | c.642_651del[ACTCATCTTG] |
| | 25 | acrR | c.-4819T>C |
| | 32 | relA | c.632_641del[ACAAACGAAT] |
| | 57 | yafW | c.150C>A |
| | 16 | metN | c.-1096C>T |
| M5 | 22 | pyrE | c.-3064del[G] |
| | 23 | acrR | c.-4819T>C |
| | 76 | yafW | c.150C>A |
| | 17 | metN | c.-1096C>T |
| | 105 | aceE | c.1874T>G |
| M6 | 35 | pyrE | c.-3064del[G] |
| | 15 | gor | c.1215T>G |
| | 20 | acrR | c.-4819T>C |
| | 15 | rhsC_3 | c.724T>G |
| | 83 | yafW | c.150C>A |
| | 32 | nanR_2 | c.-706C>T |
| | 18 | pstB | c.-148T>A |
